# Cycles in Seizure Duration and Their Underlying Dynamics in the Tetanus Toxin Rat Model

**DOI:** 10.1101/2024.11.27.625789

**Authors:** Parvin Zarei Eskikand, Sepehr Kazemi, Mark J. Cook, Anthony N. Burkitt, David B. Grayden

## Abstract

Seizure duration, a characteristic of epilepsy that is understudied in relation in its relationship with rhythmic cycles, provides critical insights into the severity and temporal dynamics of seizures. This study investigates the rhythmic patterns of seizure duration in the Tetanus Toxin (TT) rat model of epilepsy, which is a well-established platform enabling long-term, stable recordings and observation of seizure emergence and remission. Our analysis shows significant cyclical patterns in seizure durations, with periods ranging from 4 to 8 days across rats. The synchronization index and circular-linear correlations revealed phase-locked relationships between seizure durations and cycles, suggesting non-random, predictable temporal dynamics. Further analyses examined the relationship between seizure durations, inter-seizure intervals, and dominant EEG power. The relationship between inter-seizure intervals and seizure duration was modest, suggesting little to no temporal dependency. In contrast, seizure duration showed stronger associations with EEG power in dominant frequency bands, potentially reflecting fluctuations in neural excitability. The findings highlight that seizure durations exhibit predictable rhythms, which could transform seizure prediction and enable time-based intervention strategies, ultimately improving epilepsy management and patient outcomes. These insights lay the groundwork for personalized, rhythm-aware therapeutic approaches.

## Introduction

Epilepsy, a neurological disorder characterized by recurrent, unprovoked seizures, affects millions worldwide. These seizures vary significantly in frequency, duration, and severity, creating challenges for effective management and treatment. A critical aspect of epilepsy care is the ability to predict seizures accurately, as reliable forecasting could vastly improve patient safety and quality of life. Traditionally, research has focused on identifying patterns in seizure frequency [3, 4, 6, 13, 17, 18, 19, 20, 24]. However, duration patterns, which can reveal more about the intensity and potential impact of seizures, remain underexplored.

While seizure duration has historically received less attention compared to seizure frequency, several studies have examined its characteristics and relationship to seizure timing. For instance, [13] investigated correlations between seizure duration and inter-seizure intervals, suggesting that duration may be temporally structured. [7] further demonstrated that seizure durations often cluster into distinct subpopulations, implying underlying biological mechanisms that govern seizure length. Additionally, [16] examined rhythmicity in seizure occurrence and interictal spike rates over circadian and multiday timescales, underscoring the broader importance of temporal structure in epilepsy. While these studies highlight key aspects of seizure timing and variability, few have systematically explored whether seizure durations themselves follow long-term rhythmic patterns, such as circadian, multidien, or phase-locked cycles. Our study builds on this foundation by focusing specifically on the temporal evolution and cyclicity of seizure duration, aiming to uncover whether duration exhibits stable, biologically driven rhythmic patterns that could enhance personalized prediction and treatment strategies. This novel approach is grounded in the understanding that temporal dynamics—such as circadian and multi-days rhythms—affect various biological processes and may influence seizure characteristics, including duration. Detecting cyclical patterns in seizure duration could lead to breakthroughs in seizure prediction and time-based intervention strategies, improving outcomes for individuals with epilepsy. Also, understanding the temporal dynamics of seizures provides valuable insights into the underlying neurophysiological mechanisms associated with epileptic events. Due to the complexity of seizure dynamics within and across subjects, detailed longitudinal studies are essential to unveil these potential rhythmic influences.

The Tetanus Toxin (TT) rat model, which mimics human focal epilepsy by inducing spontaneous seizures following an intra-hippocampal injection of tetanus toxin, provides a suitable environment for exploring these temporal dynamics [8]. This model enables continuous, long-term monitoring of seizure activity, making it particularly useful for identifying multiday cycles and other extended patterns. This model offers a unique opportunity to study the processes underlying the development of seizures, providing insights into how changes in neural circuits over time may lead to epileptic activity. Unlike models such as kainic acid or pilocarpine, which induce chronic epilepsy without recovery, the TT model enables observation of both seizure emergence and remission. This tractable progression makes it particularly valuable for examining dynamic shifts in seizure patterns. While seizures in this model often lack a strong behavioural correlate, the ability to perform stable, long-term electroencephalogram (EEG) recordings without motion artifacts makes it ideal for analysing seizure rhythms. In this study, we leveraged the TT rat model to analyze seizure duration data collected over a 40-day period, using sinusoidal curve fitting to reveal any underlying rhythmic fluctuations. By calculating the Synchronization Index (SI) and assessing circular-linear correlations, we examined the phase-locked relationship between seizure durations and identified cycles, hypothesizing that seizure duration might exhibit phase-locked behavior indicative of intrinsic neural rhythms.

Beyond rhythmic evolution, this study explored additional factors that might affect seizure duration, specifically inter-seizure intervals and dominant EEG power. By investigating the relationship between seizure duration and the time intervals preceding or following each seizure, we aimed to determine whether longer or shorter inter-seizure intervals might impact seizure duration, offering insights into the temporal dependencies in seizure dynamics. Additionally, we assessed the correlation between seizure duration and the power of the dominant EEG frequency. The role of EEG power, particularly dominant frequency, in influencing seizure duration has been highlighted in this study and may reflect neural excitability or synchronization during seizures. Lastly, we considered the impact of external stimuli, specifically low-intensity electrical stimulation or “probing,” on seizure dynamics. Probing is commonly used in neurophysiological studies to assess cortical excitability and network connectivity. While probing is generally considered safe, this research has shown that repetitive stimulation may lower seizure thresholds or extend seizure duration. We analyzed seizure patterns during both probing-on and probing-off periods to explore how probing might modulate seizure frequency and intensity.

As the first study to investigate rhythmic evolution in seizure duration, our findings could offer new insights into the mechanisms that govern seizure dynamics. By characterizing both natural rhythmic patterns and the effects of experimental probing, we aim to contribute to a more comprehensive understanding of seizure activity and advance time-based approaches for managing epilepsy. This research may have implications for improved seizure forecasting and the development of optimized, individualized treatment strategies in epilepsy care.

## Materials and methods

### Data

The dataset analyzed in this study was derived from an intra-hippocampal tetanus toxin (TT) rat model of epilepsy, as described by Crisp et al. (2020) [8]. Seven adult male Sprague-Dawley rats (rats T1-T7) received intra-hippocampal injections of TT to induce epilepsy; rat T1 died on Day 26 and is not considered in this study. In addition to the rats in the epilepsy group, four control rats were also administered phosphate-buffered saline, but they were not included in the analysis. At the time of surgery, their body weights ranged between 250 and 370 grams. Each animal was housed separately in acrylic cages with unrestricted access to food and water. The environmental conditions in the recording room were strictly regulated, with temperatures maintained between 21°C and 26°C. A light/dark cycle of 16 hours of light (approximately 16:15–08:08 AEST) and 8 hours of darkness (08:08–16:15 AEST) was implemented using an automated timer.

Before undergoing surgery, the rats were anesthetized with a combination of Ketamine (75 mg/kg) and Xylazine (10 mg/kg). Each rat had five stainless steel electrodes implanted in the skull to enable intracranial EEG recordings. The electrode placements, referenced from the bregma (0,0), were as follows (in mm): B (1.2, +3.0), W (6.8, +3.0), G (1.2, −3.0), R (6.8, −3.0), and Ref (10.6, −3.0). The TT injection was administered stereotaxically in a volume of 300 nanoliters, delivering a dose of 30 nanograms to the epilepsy model group. The injection was targeted at the CA3 region of the hippocampus, with stereotaxic coordinates of 3.5 mm posterior to bregma, 3.0 mm lateral to the midline (left hemisphere), and 3.1 mm below the dura mater. Approximately 1-2 weeks following TT administration, rats in the epilepsy model group began exhibiting spontaneous seizures.

Spontaneous seizures were identified through EEG recordings analyzed with a custom-designed algorithm. To minimize noise and artifacts, EEG signals were processed using a high-pass filter with a 1 Hz cutoff frequency. The power of the EEG signal was computed within the 13–48 Hz frequency range, as this band is known to be relevant for detecting epileptiform activitycheung2017tracking. A seizure candidate was flagged if the instantaneous band power exceeded 3 times the 5-minute moving average for a duration of at least 9 s or exceeded seven times for at least 3 s. These thresholds were chosen to optimize the balance between sensitivity and specificity in seizure detection. To confirm a seizure, the event had to be concurrently detected in at least two electrodes, thereby reducing false positives due to localized noise or artifacts. Seizure onset was defined as the moment when the band power first exceeded the threshold, while termination was marked when it returned to or below the moving average. These strict criteria ensured accurate identification of spontaneous seizures in the EEG data [5, 8]. The seizures analyzed in this study were electrographic in nature, without an associated behavioral component.

To reduce false positives from noise or local artifacts, seizure events were only included if they were detected concurrently on at least two intracranial electrodes. While this requirement may bias detection toward more spatially distributed seizures, it ensures high specificity and consistency in seizure identification. The dual-threshold algorithm used in this study was previously validated in the context of this model [5, 8] and has been shown to capture rhythmic seizure dynamics reliably. To further support transparency, example EEG traces of detected seizure events have been provided in the Supplementary Material (Fig. S1). These examples illustrate the electrographic patterns that meet detection criteria and help clarify how the algorithm distinguishes seizures from background activity or artifacts.

Seizure frequency varied daily over a period of approximately six weeks. Around 4 to 5 weeks post-injection, the frequency of seizures began to decline, eventually leading to the cessation of detectable seizure activity. EEG recordings were collected continuously at a sampling rate of 2048 Hz for 23 hours each day, with a 1-hour break allocated for maintenance and data backup.

The experimental protocol included periodic, low-level electrical stimulation or “probing” [8]. The probing-on period consisted of 100 biphasic pulses (pulse-width 0.5 ms and current/phase 1.26 *±* 0.024 mA) with inter-stimulus intervals of 3.01 s over a period of about 5 minutes. After each probing-on period, there was a 5 min probing-off period before the start of the next probing-on period. The research procedures received approval from the St. Vincent’s Hospital, Melbourne Animal Ethics Committee and were carried out following the guidelines outlined in the “Australian Code for the Care and Use of Animals for Scientific Purposes, 8th Edition” (2013).

Fig 1A displays a histogram of seizure durations across different rats, highlighting the distribution of seizure durations for each individual rat. Fig 1B displays the ranges number of seizures across different rats over 40 days of recording compared to the average duration of seizures that occurred on each day across all rats. This figure shows the daily averages across all rats, providing a comparative view of seizure frequency and duration trends over time.

**Fig 1.**
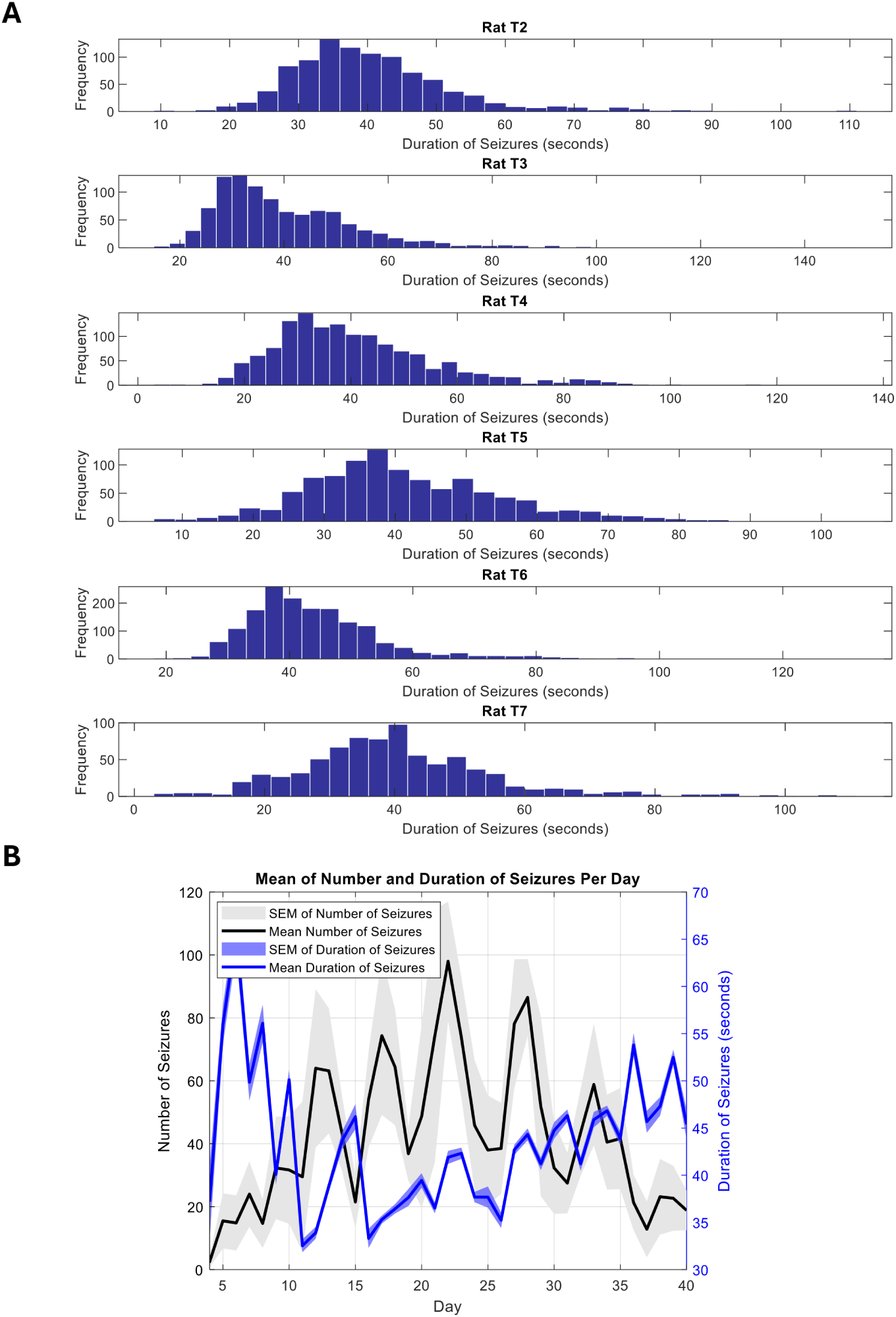
**A) Histogram of seizure durations across six rats, showing the frequency of seizure durations regardless of whether they occurred during on or off periods.** The x-axis denotes the duration of seizures in seconds, while the y-axis represents the frequency of seizures within each duration bin. **B) The mean number and duration of seizures per day over a 40-day period.** The black line represents the mean number of seizures per day, with the corresponding shaded gray area indicating the standard error of the mean (SEM) for seizure counts. The blue line represents the mean duration of seizures (in seconds), with the shaded blue area representing the SEM of seizure duration. The left y-axis corresponds to the number of seizures, while the right y-axis corresponds to seizure duration in seconds.

### Temporal analysis of the changes in the duration of seizures

We calculated the hourly average of seizure durations over 40 days of recording across multiple rats and analyzed this data to identify the underlying periodic patterns in the changes in seizure duration using sinusoidal curve fitting. The sinusoidal model applied is expressed as

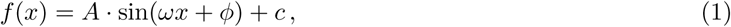

where *A* represents the amplitude, *ω* is the angular frequency, *ϕ* denotes the phase shift, and *c* is the vertical offset. The optimization process involved minimizing the Sum of Squared Residuals (SSR) between the measured data and the fitted sinusoidal function. The periods corresponding to each rat were determined and documented.

We calculated the Synchronization Index (SI) for the significant periods identified by sinusoidal curve fitting. For each rat, the positional phases of seizures within a cycle were calculated by mapping seizure occurrences to a sinusoidal cycle. To calculate the Synchronization Index (*SI*), we first converted the seizure phases to their complex exponential representation [20]. The mean resultant vector was computed as

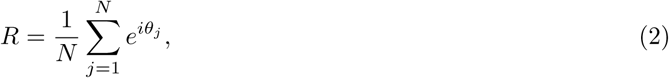

where *θ_j_* represents the phase of each seizure and *N* is the total number of seizures. The SI is then defined as the magnitude of the mean resultant vector,

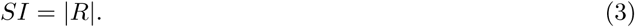

A higher SI value (closer to 1) indicates a stronger synchronization of seizures within the periodic cycle, while a lower value (closer to 0) indicates less synchronization. We reported the SI for each rat, along with the identified periods from the sinusoidal fits as performed by [20].

To assess the relationship between seizure phases within cycles and seizure durations, we calculated the circular-linear correlation [21]. In this analysis, seizure phases represent circular data, while seizure durations are linear data. The circular-linear correlation coefficient is computed as

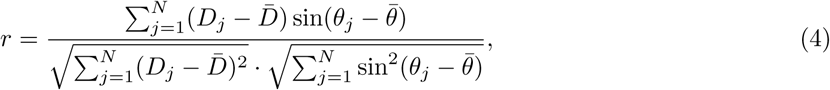

where *D_j_* is the duration of the *j*-th seizure, *θ_j_* is its corresponding phase, 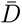 is the mean seizure duration, and 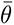 is the mean phase. This correlation coefficient quantifies the degree of association between seizure phases within cycle and duration of seizures, ranging from *−*1 (perfect negative correlation) to 1 (perfect positive correlation), with 0 indicating no correlation.

To evaluate the statistical significance of the circular-linear correlation, we performed a permutation test. This test assesses whether the observed correlation could have arisen by chance. We calculated the observed circular-linear correlation coefficient, *r*_obs_, using the actual data. We then randomly shuffled the seizure durations while keeping the phases unchanged. For each permutation, we recalculated the circular-linear correlation coefficient, generating a distribution of correlation coefficients under the null hypothesis (i.e., that there is no relationship between seizure phases and durations. This process was repeated 100,000 times to build a distribution of permuted correlation coefficients. The p-value was determined by comparing *r*_obs_ to the distribution of permuted correlations*r*_per_. Specifically, the p-value was calculated as the proportion of permuted correlations that were as extreme as or more extreme than the observed correlation,

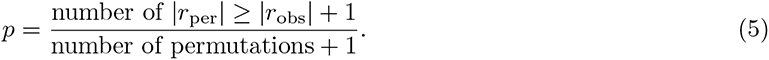

To correct for multiple comparisons across rats, the calculated p-value was multiplied by the number of comparisons (i.e., the number of rats), ensuring that the results were adjusted for family-wise error rates. The permutation test ensures a robust assessment of the statistical significance of the circular-linear correlation, with p-values less than 0.05 considered significant. As a final step, the data was divided into 15-day segments (360 hours each) with a 359-hour overlap to better demonstrate changes over time on the cycles as the seizures waxed and waned. Periodic patterns were analyzed over time, and significant cycle within each segment were identified using the method described above.

It is important to note that the Synchronization Index (SI) quantifies the concentration or clustering of seizure events (or seizure durations) around a specific phase of a cycle. In our context, a high SI indicates that seizure durations tend to occur more consistently at a particular phase of the multiday rhythm, regardless of their actual value. In contrast, the circular-linear correlation measures the strength and direction of association between the phase of the cycle (circular variable) and the seizure duration (linear variable).

### Is seizure duration determined by the cycle or the time interval between seizures?

We investigated the correlation between the time before a seizure and the duration of the seizure. Additionally, we analyzed whether seizure duration influences the time until the next seizure. Beyond these analyses, we conducted a survival analysis to explore whether the time before a seizure affects the probability of the next seizure occurring (in line with the classical *seizure begets seizure* theory [15]) and whether seizure duration influences the likelihood of a subsequent seizure.

Survival analysis was performed using seizure data from six rats (T2-T7). Two primary variables were used for the analysis: seizure duration and time before seizures (time elapsed since the previous seizure). For each rat, the time to the next seizure was used as the survival time. The survival time was defined as the time between consecutive seizures, while the time before each seizure and duration of seizure were used as two covariates. A Cox proportional hazards model was fitted separately for each rat using the seizure duration and time before seizure as covariates [14]. The coxphfit function in MATLAB was used to fit the model, with the formula,

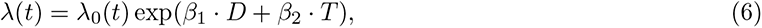

where *D* is the duration of the seizure, *T* is the time before the seizure, *λ*(*t*) is the hazard function, *λ*_0_(*t*) is the baseline hazard, *β*_1_ is the coefficient for seizure duration, and *β*_2_ is the coefficient for the time before the seizure.

The values of *β*_1_ and *β*_2_ are estimated from the data using maximum likelihood estimation. The hazard ratios (HR) were calculated by exponentiating the model coefficients, and 95% confidence intervals (CI) were computed for the hazard ratios using the standard errors from the model.

### The correlation between dominant frequency power and seizure duration

We also explored whether there is a correlation between the power in the dominant frequency and the duration of seizures. This analysis provides insights into the factors that influence seizure duration and examines the possibility of cycles in EEG power at certain frequencies. To analyze the dominant frequency of the EEG during seizures in probing off periods, logarithmic frequency bands were defined from 1 Hz to 130 Hz. A total of 10 logarithmically spaced frequency bands were created to capture the frequency content of the EEG. For each seizure, the power spectral density (PSD) was computed using the Welch method [23]. The dominant frequency for each seizure was defined as the frequency band with the maximum power. Dominant EEG power was computed during seizure events only, using data from probing-off periods to avoid contamination from stimulation artifacts.

### Statistical analysis

Different statistical tests were applied depending on the nature of the hypothesis and data structure to assess rhythmicity, temporal associations, and group differences in seizure duration across multiple conditions. To evaluate phase-locking of seizure durations to identified cycles, the Synchronization Index (SI) was calculated, and circular-linear correlations were computed to quantify the association between seizure duration (a linear variable) and seizure phase (a circular variable). The statistical significance of the circular-linear correlations was determined using a non-parametric permutation test with 100,000 iterations, and p-values were adjusted for multiple comparisons using the Bonferroni correction. To examine the relationship between seizure duration and inter-seizure intervals, a Cox proportional hazards model was applied, using seizure duration and time since the previous seizure as covariates. To assess differences in seizure durations between probing-on and probing-off periods, Welch’s two-sample t-tests were used, which account for unequal variances between groups. Additionally, Pearson correlation coefficients were calculated to evaluate the linear relationship between seizure duration and power in the dominant EEG frequency band during probing-off periods. For all statistical tests, a significance threshold of *p <* 0.05 was used.

## Results

### Periodic patterns and phase-locked rhythms in seizure durations

The analysis of seizure data for six rats over a period of 40 days revealed distinct periodic patterns for seizure duration of each rat. Fig 2A shows the raster plot of the seizure duration and the best fitted sinusoid to the data. We assumed that the cycle frequencies were fixed over the whole recording periods. The results demonstrate clear cyclical patterns in seizure durations across all six rats (T2–T7) over the 40-day recording period. For each rat, the average hourly seizure durations (blue dots) fluctuate over time, and the fitted sinusoidal curves (cyan lines) capture these periodic changes. The periods of the fitted sinusoidal functions vary across the rats, ranging from 4.1 days (Rat T2) to 8.6 days (Rat T5), indicating individual variability in the seizure cycles. The fitted sinusoidal curves highlight the presence of rhythmic fluctuations in seizure durations, suggesting a temporal structure in seizure occurrence that may be tied to underlying physiological or environmental rhythms.

**Fig 2.**
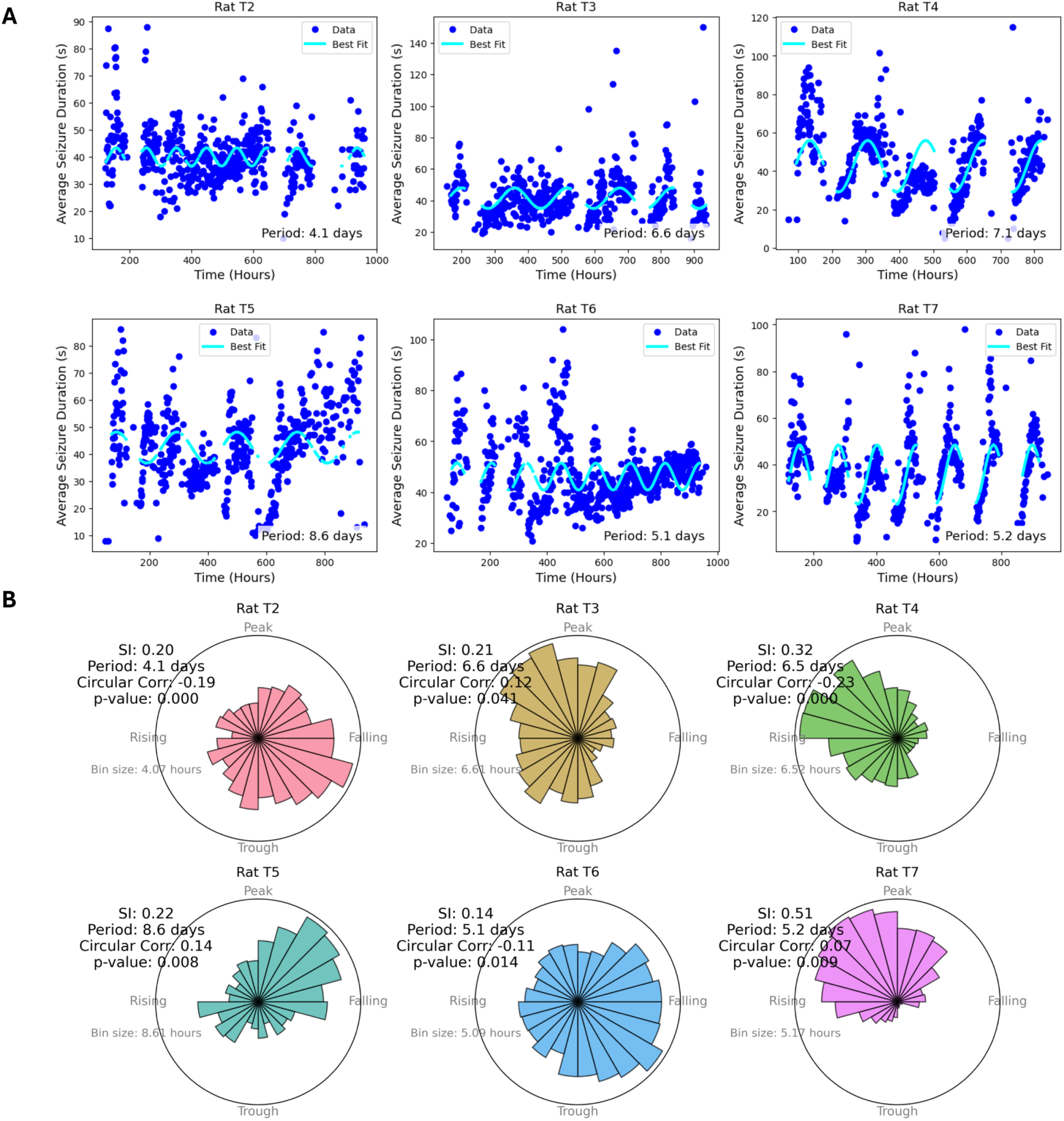
A) Raster plots of seizure durations over 40 days (960 hours) of recording for six rats (T2–T7). Each blue dot represents the average seizure duration recorded over a one-hour interval. The cyan curve represents the best-fit sinusoidal function to the data, capturing periodic patterns in seizure durations. The period of the fitted sinusoidal is indicated for each rat, revealing distinct cyclical patterns in seizure durations. **B) Polar histograms displaying the relationship between seizure durations and the phases of the identified cycles.** Each subplot represents a different rat (T2 through T7). Each radial bar represents the distribution of seizure duration within different phases of the cycle. The predominant period and the Synchronization Index (SI), Eq. (3), are annotated on each plot, with higher SI values indicating stronger phase locking. Circular-linear correlations between the phase of the cycles and seizure durations are provided. Reported *p*-values represent the proportion of permuted correlations exceeding the observed correlation, corrected for multiple comparisons using the Bonferroni method. Significant *p*-values indicate that seizure durations are non-randomly distributed across the cycle phase in most rats.

Fig 2B presents the polar histograms of the relationship between seizure durations and the phases of the identified cycles. The results indicate a clear relationship between the phase of the identified cycles and the durations of seizures across all six rats (T2–T7). The clustering of durations of seizures in a specific phase interval implies that seizure durations are not random but follow a predictable pattern. The Synchronization Index (SI) values, which range from 0.14 to 0.51, indicate varying degrees of phase locking, with higher SI values reflecting stronger clustering of seizures at specific phases of the cycle. Rats T4 and T7 exhibit the highest SI values (0.32 and 0.51, respectively), implying a stronger synchronization between seizure durations and the cycle phases. In contrast, Rat T6 shows the weakest phase locking, with an SI value of 0.14. The circular-linear correlations between the phase of the cycles and seizure durations are significant in most rats, as evidenced by the low p-values. These significant correlations suggest that seizure durations are non-randomly distributed across different phases of the cycle, indicating that seizures tend to occur more frequently or with longer durations during certain phases of the cycle. These results highlight a cyclical, rhythmic influence on seizure durations, with substantial individual variability in the strength of phase locking across the rats.

The Tetanus Toxin model of epilepsy is characterized by fluctuating activity over the weeks of recording, with seizure properties evolving over time [8]. To examine these temporal changes, the data was divided into 360-hour (15-day) segments, with each segment advancing by 1 hour. This method enabled the analysis of seizure cycles across various time windows during the 40-day recording period. A sinusoidal curve was fitted to the data for each segment to capture periodic patterns. Fig 3 visualizes the temporal evolution of seizure durations across individual rats, showing variability in the strength and consistency of rhythmic patterns. The results reveal dynamic changes in seizure cycles over time for all six rats (T2–T7). In each subplot, significant periods of cyclical seizure durations are identified. These periods are represented by horizontal lines, with their color indicating the significant of correlations between seizure phases and duration of seizures across overlapping 15-day segments. The figure reveals that the periodicity of seizure durations changes over time, with different rats exhibiting varying patterns and strengths of significant cycles across the recording period. Notably, rat T7 demonstrates a clear and persistent cyclical trend in seizure duration over time, while other animals (e.g., T2, T3) show more variable or intermittent patterns. This figure supports the conclusion that seizure duration rhythmicity is subject-specific, and highlights the unique stability observed in T7. Such variation may reflect underlying biological or network-level differences between animals. The presence of strong correlations (low p-values) suggests that seizure durations follow rhythmic, cyclical patterns during these periods.

**Fig 3.**
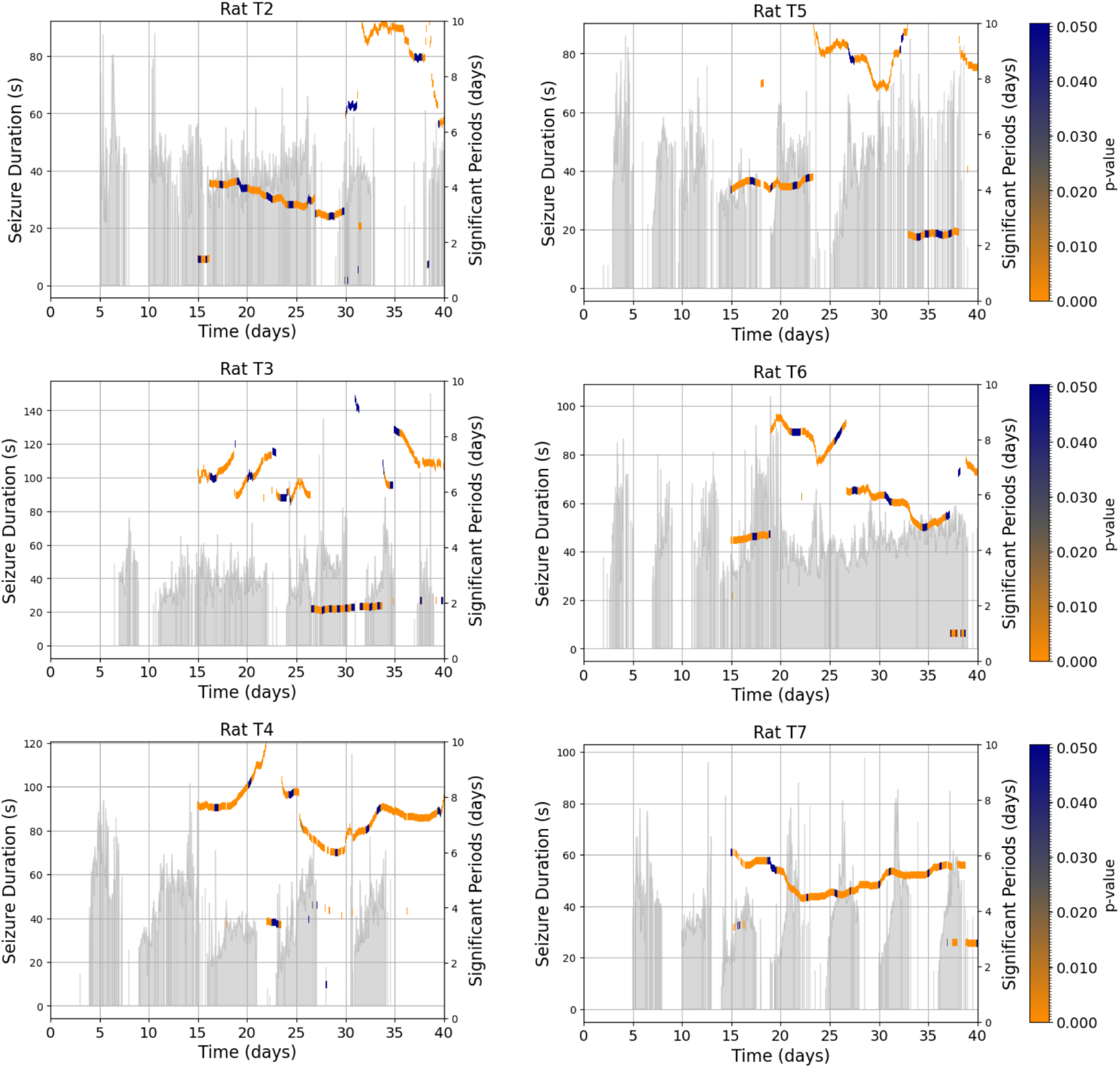
Changes in seizure cycles over time for six rats (T2–T7). Each subplot displays the significant periods (in days) of cyclical changes in seizure durations as horizontal lines, plotted against time (in days) across multiple overlapping segments. Each segment spans 360 hours (15 days) and overlaps with the next by 359 hours to enable continuous tracking of rhythmicity. The color of each line represents the p-value of the correlation between seizure duration and the phase of the detected cycle (color bar on right). P-values were derived from a nonparametric permutation test assessing the strength of circular-linear correlations between seizure duration and seizure phase. Low p-values (stronger phase-locking) are shown in purple, while higher p-values are in blue. Gray histograms on the left axis of each subplot show hourly seizure durations over the 40-day recording window. This figure illustrates subject-specific variability in rhythmic seizure duration patterns: Rat T7 shows a persistent and robust rhythmic structure, while other animals exhibit more variable or weaker phase relationships. These differences highlight biological heterogeneity in rhythmic expression across subjects.

### Is there a relationship between the time interval between seizures and the duration of seizures?

The results suggest that there is a relationship between the time intervals separating seizures (only in extreme cases where the inter-ictal interval is either very short or very long) and the durations of seizures. Extreme cases were selected for this analysis, comparing very long inter-ictal intervals (greater than or equal to 1 day) and very short inter-ictal intervals (less than 1.5 minutes) between seizures. TTo assess whether seizure duration was influenced by the temporal proximity of prior events, we compared seizures that occurred after long inter-seizure intervals (*≥*1 day) with those that followed shortly after a previous seizure (*<*1.5 minutes). These threshold values were chosen to contrast two extreme recovery scenarios—prolonged versus immediate recurrence, while ensuring that each group retained a sufficient number of seizures for statistical analysis (e.g., 23–27 events per group). While the sample size is modest, this approach allows us to probe possible effects of inter-event timing on seizure characteristics.

Seizures that occur after a long inter-ictal interval (*≥* 1 day) tend to have shorter durations, with a median seizure duration of approximately 20-25 seconds. When the next seizure occurs after a long inter-ictal interval (*≥* 1 day), the seizure duration appear to be longer, with a median duration of around 40-45 seconds. Seizures occurring after a very short inter-ictal interval (less than 1.5 minutes) are relatively shorter, with median durations around 35 seconds (Fig 4A).

**Fig 4.**
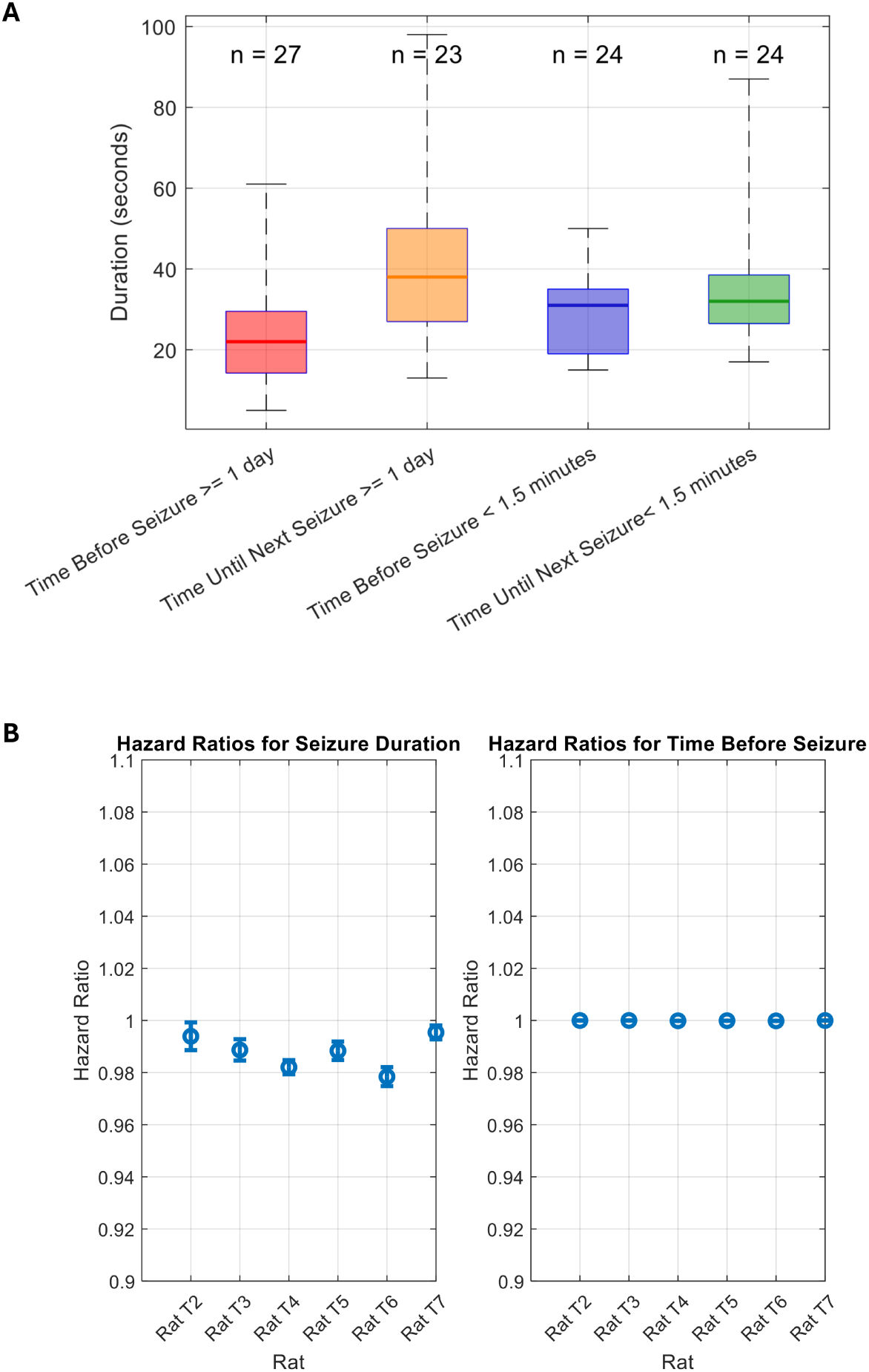
A) Boxplot illustrating the relationship between the time interval separating seizures and the duration of seizures. Four extreme conditions are shown: time before the previous seizure is greater than 1 day, time until the next seizure is greater than 1 day, time before the previous seizure is less than 1.5 minutes, and time until the next seizure is less than 1.5 minutes. Each box represents the distribution of seizure durations (in seconds) for the respective condition, with the horizontal line in the box indicating the median duration. The number of seizures (*n*) for each condition is displayed above the box. This comparison explores whether the time interval between seizures influences the duration of the subsequent seizure. **B) Hazard ratios for seizure duration and time before seizure across rats.** The plot on the left shows the hazard ratios for the effect of seizure duration on the risk of subsequent seizures in each rat, with confidence intervals. The plot on the right shows the hazard ratios for the effect of time before the previous seizure on the risk of subsequent seizures. Data is presented for rats T2 through T7, with hazard ratios below 1 indicating a reduced risk of seizure occurrence as the covariates increase. Error bars represent 95% confidence intervals.

The results of the Cox proportional hazards model analysis for seizure duration and time before seizure across six rats (T2 through T7) are shown in Fig 4B. The left panel presents the hazard ratios for seizure duration, indicating if longer seizure durations are associated with a reduced risk of subsequent seizures. The hazard ratios for seizure duration across all rats are slightly below 1, suggesting that an increase in seizure duration is associated with a modestly reduced risk of a subsequent seizure. In the right panel, the hazard ratios for the time before the previous seizure are depicted. These hazard ratios are close to 1 but exhibit some variability across rats. For most rats, the time before the previous seizure has no effect on the risk of the next seizure.The effect sizes, reflected in hazard ratios, were modest across animals. In both analyses, the hazard ratios remain close to 1, indicating that neither seizure duration nor time before seizure has a strong influence on the likelihood of subsequent seizures.

To further explore how long seizure-free intervals influence the timing of subsequent seizures, we examined the inter-seizure interval following seizures that occurred after a delay of *≥*1 day. As shown in Fig. S2, these events were typically followed by much shorter inter-seizure intervals (median *<* 5 hours), indicating that a long recovery period is often followed by a rapid recurrence. This supports the idea that seizures emerging after prolonged quiescence may initiate a cluster or or occur within a phase of increased seizure susceptibility.

### Dominant frequency and the duration of seizures

We investigated whether there is a correlation between seizure duration and the power of the dominant EEG frequency during the probing off period. The results, displayed in Fig 5A, indicate that there is a statistically significant correlation between these two variables across all six rats (T2 to T7). All rats exhibit moderate to strong positive correlations between seizure duration and dominant EEG power. This suggests that, for these rats, longer seizures tend to be associated with higher dominant EEG power.

**Fig 5.**
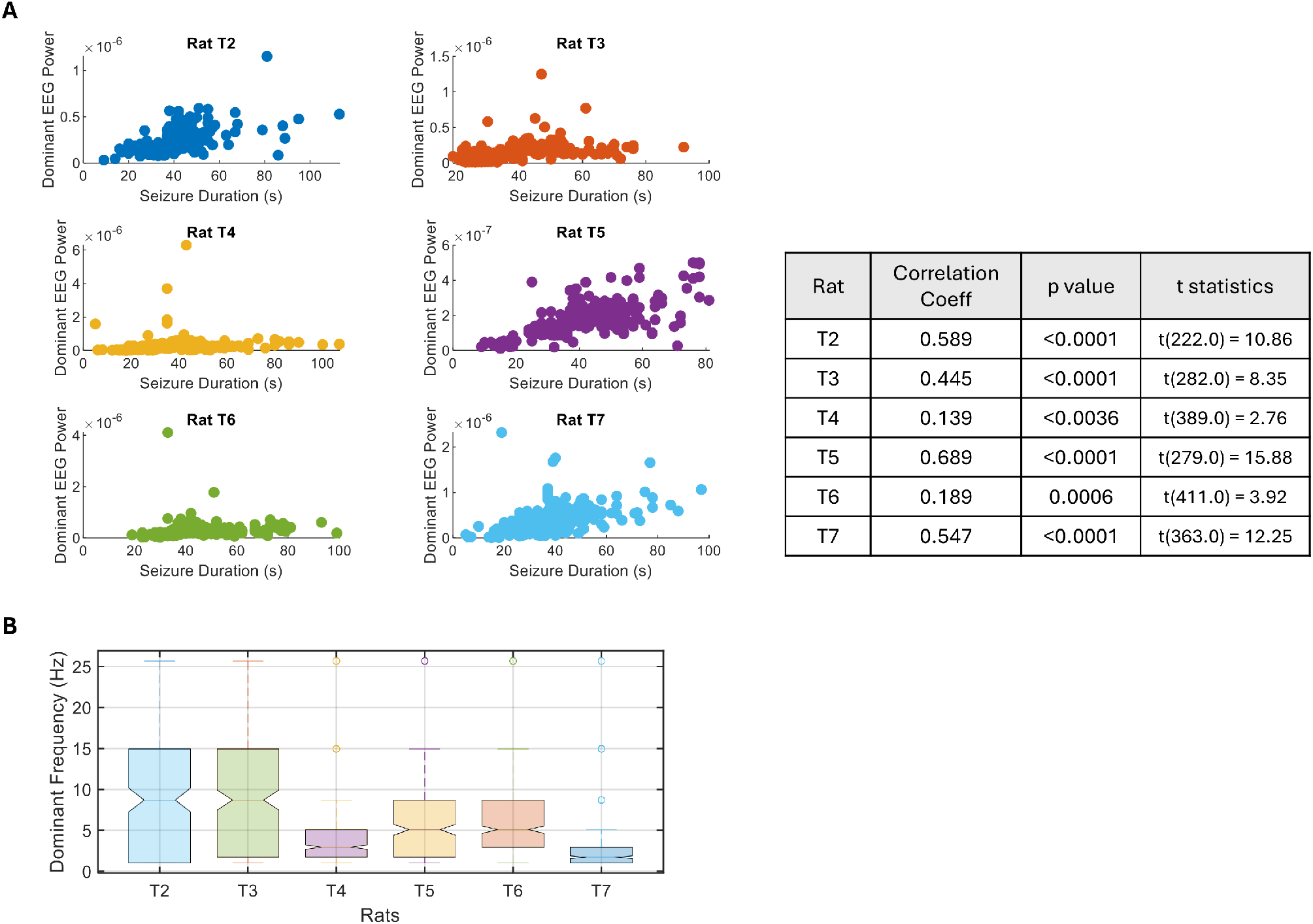
Relationship between seizure duration and power of dominant frequency during the probing off period. A) Scatter plots display the correlation between seizure duration and power of dominant frequency for six rats (T2 to T7). Each plot represents the individual relationship between seizure duration and the dominant EEG power recorded during that seizure. The table on the right summarizes the Pearson correlation coefficients, corresponding p-values, and associated *t*-statistics for each rat. B) Distribution of dominant EEG frequencies for all rats. The notched box plots display the distribution of the dominant EEG frequencies (in Hz) for each rat (T2 through T7) during seizures. The notches indicate the confidence interval around the median, and the whiskers show the range of the data. Outliers are represented by individual points beyond the whiskers. These dominant frequencies were found to have strong correlations with seizure duration, as shown in Fig 5A.

Looking at the distribution of dominant frequencies for each rat (T2 through T7) at Fig 5B, we can observe the range of frequencies in relation to known EEG frequency bands. The dominant frequencies in rat T2 are mostly distributed in the alpha range, with the majority of the data clustered between 5-15 Hz. This indicates that many seizures in this rat are associated with activity in the alpha band. Similar to T2, Rat T3 exhibits a broad distribution of frequencies, but with a slightly higher range, including both alpha and low beta frequencies (13-20 Hz). There is a greater spread in the data, though the frequencies still tend to be within or just above the alpha range. The dominant frequencies for Rat T4 are mostly concentrated in the delta range (1-4 Hz), with very few instances exceeding the alpha range. This suggests that seizure activity in Rat T4 is more likely associated with lower-frequency EEG activity, potentially indicating different neural dynamics compared to other rats. The distribution of frequencies for Rat T5 is extending well into the alpha range and higher. While some frequencies fall into the theta (4-8 Hz) and delta ranges, there are significant occurrences of dominant frequencies in the alpha range, similar to Rats T2 and T3. The frequencies for Rat T6 are primarily concentrated in the alpha range (8-12 Hz) and theta range (4-8 Hz). The distribution for Rat T7 is focused at the lower end of the spectrum, with many values in the delta range, similar to Rat T4. This suggests seizure activity that is more associated with low-frequency oscillations.

### Seizure durations during probing on periods compared to probing off periods

Fig 6 displays a histogram of seizure durations across different rats that are occurred during the probing on periods compared to probing off periods. The results indicate notable differences in the distribution of seizure durations between the probing on and off periods across all six rats (T2–T7). The histograms show a higher frequency of seizures during the on periods compared to the off. The results in Fig 6b show distinct differences in seizure durations between probing on periods and probing off periods for several rats. Rats T2, T3, T4, and T7 exhibit significant differences in seizure durations, as indicated by p-values below 0.05. In these cases, seizures during the on periods tend to have longer durations compared to those during off periods.

**Fig 6.**
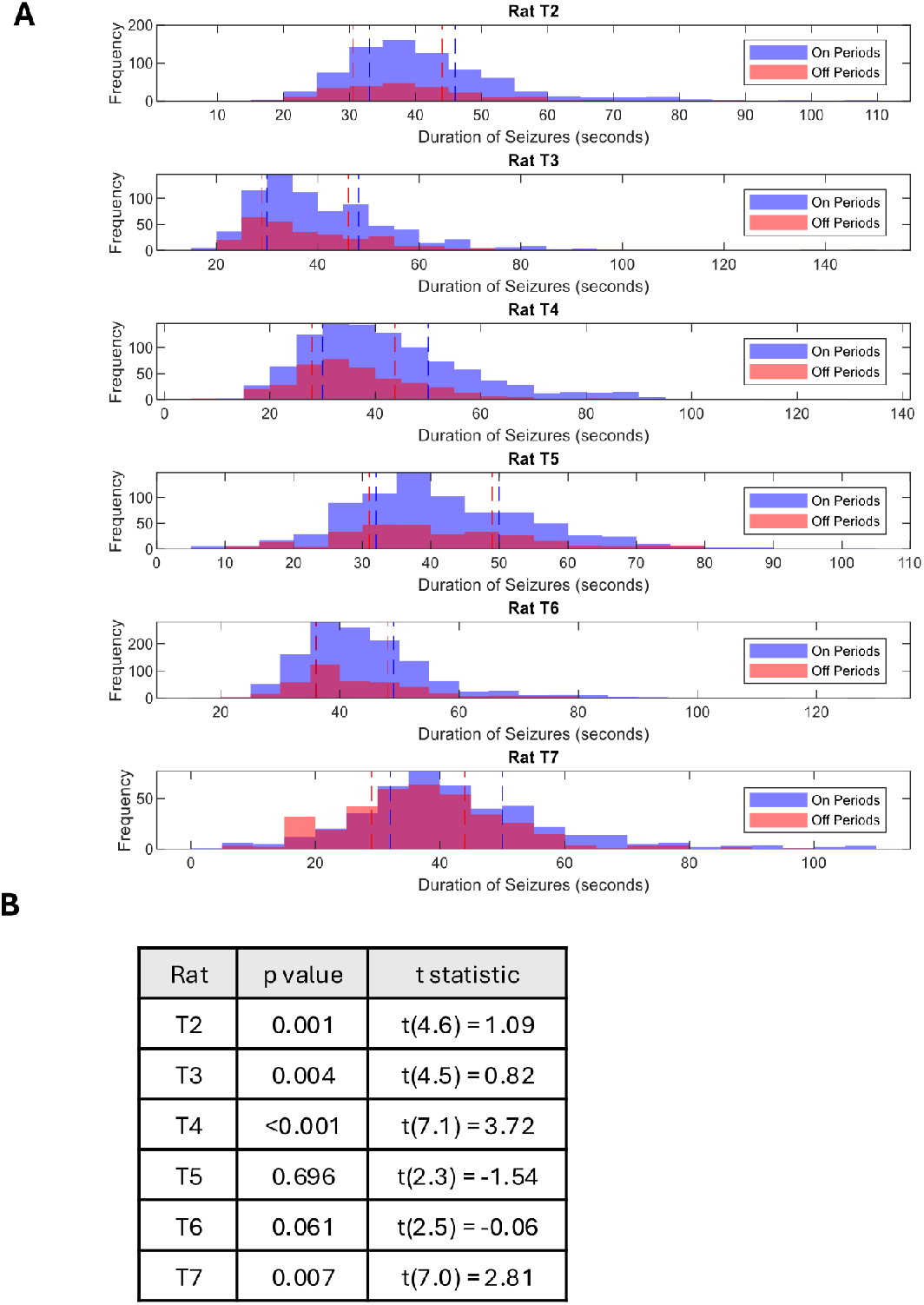
Comparing seizure durations during the on and off periods. a) The histograms of seizure duration during the on and off periods.The blue bars represent the distribution of seizure durations during the probing on periods, while the red bars represent those during the probing off periods. Dashed vertical lines indicate the low (left) and high (right) quantiles for each period. b) Welch’s two-sample t-test was used to compare seizure durations between on and off periods for each rat. The table reports the t-statistics, degrees of freedom, and corresponding p-values. Significant differences (p ¡ 0.05) were observed in most rats (T2, T3, T4, and T7), as indicated by p-values less than 0.05.

## Data availability statement

Data are available upon reasonable request from the corresponding authors. Code is available on Github (https://github.com/ParvinZE/Cycles-in-Seizure-Duration).

## Discussion

In this study, we explored the factors that could influence the duration of seizures, aiming to better understand what controls the severity of seizures. Specifically, we investigated the presence of rhythmic patterns in seizure duration and examined whether the interval between seizures affects the duration of the next seizure. Through a comprehensive analysis of seizure data from six rats, we identified several key insights into the temporal dynamics of seizure duration and the potential factors contributing to its variability.

### Cyclical patterns in seizure duration

The results of this study highlight the presence of significant cyclical patterns in seizure durations across six rats (T2–T7) over the 40-day recording period. The sinusoidal fits to the seizure duration data captured clear rhythmic fluctuations for each rat, suggesting that seizure durations follow predictable temporal structures. These findings support the idea that seizure duration may be influenced by underlying physiological or environmental rhythms that manifest as distinct cycles. While all rats exhibited rhythmic patterns, the strength and regularity of these cycles varied, with periods ranging from 4.1 days in rat T2 to 8.6 days in rat T5, highlighting the individual variability in the temporal dynamics of seizure durations. Despite these differences in periodicity, the presence of clear cyclical patterns in all six rats suggests that seizure durations are not randomly distributed but are instead governed by time-based processes.

The polar histograms and Synchronization Index (SI) analyses further supported these observations, revealing significant relationships between seizure durations and the phases of identified cycles. The clustering of seizure durations at specific phases of the cycle, particularly in rats with higher SI values (T4 and T7), suggests that seizure timing and duration are influenced by underlying cyclic rhythms. This phase locking of seizure durations reflects a non-random distribution of seizures, reinforcing the idea that seizure occurrence and duration are governed by predictable temporal dynamics. The circular-linear correlations between seizure durations and the phases of the identified cycles further reinforce the notion of non-random patterns in seizure activity. These correlations are significant in all rats, indicating that seizure durations are not uniformly distributed across the cycle phases. Instead, certain phases of the cycle correspond to longer or shorter seizure durations, suggesting a predictable relationship between seizure timing and seizure duration.

The rhythmic patterns in seizure duration observed in this study occur on multiday timescales—longer than a typical circadian rhythm. While circadian modulation of seizures is well documented, our findings align more closely with multidien patterns previously reported in [3]. These non-circadian rhythms suggest that seizure features, including duration, may be governed by slow, endogenous cycles that vary across species. Additionally, it is important to note that seizure durations were averaged over hourly bins to balance temporal resolution with analytical tractability and reduce noise from highly irregular inter-event spacing. However, we acknowledge that this approach may introduce smoothing effects that obscure faster or more transient rhythmic patterns, particularly those operating on sub-hourly timescales.

### Temporal dynamics of seizure duration

The Tetanus Toxin model of epilepsy is known for its evolving seizure activity over time. The dynamic nature of seizure cycles is further underscored by the analysis of changing seizure patterns over time. The use of overlapping time segments allowed for the detection of evolving cyclical patterns throughout the 40-day recording period. In rats like T7, where significant cycles were consistently detected, there appears to be a more stable rhythmic influence on seizure durations over time. In contrast, the more variable cycles observed in other rats highlight the dynamic nature of seizure patterns. Despite these variations, the results also reveal a general decreasing pattern in the periods of cycles during the mid-recording phase, particularly between day 20 and day 30 for Rats T2, T4, and T6. In these rats, the periods of significant seizure duration cycles become shorter over this time window, indicating a shift in the underlying rhythmic processes. The consistent and strong rhythmicity observed in rat T7 raises the possibility of underlying biological or structural differences contributing to this pattern. While we did not perform post-hoc histological analysis in the current study, variability in the extent of injury caused by tetanus toxin injection may influence the stability and expression of seizure rhythms.

Additionally, we acknowledge that neural plasticity during seizure remission and recovery may influence the temporal dynamics observed in seizure durations. This plasticity introduces a dynamic backdrop that could modulate rhythmic patterns in seizure activity. Rather than viewing this entirely as a confound, we interpret it as an opportunity to understand how seizures evolve and dissipate over time. Our rhythmic analyses were conducted with this evolving baseline in mind, and the results should be considered in the context of a plastic, recovering neural network.

We compared these results with the dynamics of the cycles in the occurrence of seizures studied Figure 6 of of [27]. There are some similarities between the evolving patterns of the periods of cycles in the number of seizures and the duration of seizures across the rats studied. For instance, in rat T2, there is a general reducing pattern in the periods of cycles from day 15 to day 30 in both the number of seizures and seizure durations. This reduction in period length across both metrics suggests the possibility of a shared underlying rhythm that affects both the frequency of seizures and their duration over this specific time window. Similarly, rat T7 shows a stable periodicity in both seizure number and duration throughout the entire 40-day period. The consistent cycles in both figures suggest a strong, predictable rhythm influencing both aspects of seizure activity, indicating that the temporal dynamics of seizure occurrence and duration are tightly linked in this rat. In rat T4, a distinct pattern emerges between day 30 and day 40, where the identified period fluctuates slowly around 7 days in both the seizure number and seizure duration.

These specific examples illustrate that the cycles governing seizure frequency and duration are synchronized. However, the degree of similarity between seizure number and duration cycles varies across the rats, indicating individual variability in the rhythmicity of seizures. This underscores the importance of further investigating these temporal patterns to improve seizure forecasting and develop personalized epilepsy management strategies.

### The relationship between seizure duration and inter-seizure intervals

The relationship between the time interval separating seizures and the duration of seizures reveals interesting patterns, particularly in extreme cases where the inter-ictal interval is either very short (less than 1.5 minutes) or very long (greater than or equal to 1 day). The results suggest a potential link between the duration of seizures and both the time before and the time until the next seizure, though this effect is only pronounced in these extreme cases. Seizures that occur after a long inter-ictal interval of greater than or equal to 1 day tend to have shorter durations, with a median duration of 20–25 seconds. This pattern suggests that longer recovery periods between seizures may result in briefer seizure events. Conversely, when seizure durations are relatively long, there is often a longer inter-ictal interval until the next seizure, with the median duration of these longer seizures being around 40–45 seconds. This relationship is particularly evident in extreme cases where the time until the next seizure exceeds 1 day, indicating that longer seizure events may be associated with greater spacing between seizures in these instances. This is in agreement with the study by Ferastraoaru et al. (2016) on humans with focal epilepsy, which showed that the terminal seizure (the last seizure in a cluster) tends to be longer, similar in duration to isolated seizures [11].

In order to further investigate the relationship between the time interval separating seizures and seizure duration, we conducted a survival analysis using the Cox proportional hazards model. This analysis allowed us to assess whether longer seizure durations or the time before the previous seizure influence the risk of a subsequent seizure. The results of this analysis, presented in Fig 4B, show the hazard ratios for both seizure duration (left panel) and the time before the previous seizure (right panel) across six rats (T2 through T7). The hazard ratios for seizure duration are all slightly below 1, suggesting that longer seizure durations are modestly associated with a reduced risk of a subsequent seizure. This pattern implies that longer seizures may provide some exhaustive effect, potentially increasing the inter-ictal interval until the next seizure. However, this effect is relatively small, as the hazard ratios remain close to 1.

Taken together, these results suggest that neither seizure duration nor the time before the previous seizure has a strong influence on the timing of subsequent seizures. While longer seizure durations may slightly reduce the risk of another seizure, the overall impact is minimal. These findings highlight the complex nature of seizure dynamics and suggest that factors other than seizure duration and interseizure intervals, such as the dynamic cycles, play a more critical role in determining seizure timing. The study by Cook et al. (2016) [7] conducted on human subjects with focal epilepsy also showed that no linear association between inter-seizure intervals and durations, suggesting that a longer buildup (inter-seizure interval) does not necessarily result in a proportionally longer seizure duration (discharge). However, in two subjects, the study was able to investigate a logistic relationship between inter-seizure intervals and durations, which revealed that short-duration seizures were more likely to follow short inter-seizure intervals, and longer inter-seizure intervals were more likely to be followed by longer seizures [7].

### Correlation between seizure duration and dominant EEG frequency: insights into neural oscillations and stability of seizure cycles

The relationship between seizure duration and the power of the dominant EEG frequency was explored to determine if longer seizures are associated with higher EEG power during the probing off period. The results, as shown in Fig 5A, reveal a statistically significant positive correlation between seizure duration and dominant EEG power across all six rats (T2 to T7). This consistent pattern of moderate to strong correlations suggests that, in these rats, longer seizures tend to be linked with increased dominant EEG power, indicating a possible interaction between seizure intensity and neural oscillations. For example, rat T5 exhibited the strongest correlation (r = 0.689), while Rat T4 showed the weakest (r = 0.139), though still statistically significant. These findings imply that seizure dynamics, particularly duration, may be closely tied to the dominant EEG power. In general, this suggests that, as seizures last longer, the power of the dominant EEG frequency during those events increases, possibly reflecting greater neural involvement or heightened brain activity during prolonged seizures.

The distribution of dominant frequencies across the six rats, displayed in Fig 5B, provides additional context to the correlations observed. The dominant frequencies for most rats tend to fall within known EEG frequency bands, though there are individual variations. For example, rats T2, T3, T5, and T6 primarily exhibit dominant frequencies in the alpha range (8-12 Hz), indicating that their seizures are associated with higher-frequency brain activity. This could reflect more synchronous neural firing patterns during these seizures. In contrast, rats T4 and T7 have dominant frequencies that are concentrated in the delta range (1-4 Hz), which is characteristic of slower, large-scale neural oscillations. Interestingly, these two rats also display more stable periods of cycles during the entire recording compared to the other rats and have a stronger Synchronization Index (SI) with the identified periods from sinusoidal fitting. This suggests a potential relationship between the dominance of delta-range frequencies and the stability of the seizure cycles. The lower-frequency oscillations in the delta range may reflect a more regular, rhythmic neural activity, which could result in more stable cyclic patterns observed in rats T4 and T7. The differences in frequency distribution between rats suggest distinct neural dynamics underlying their seizure activity. For instance, the delta-dominated seizures in rats T4 and T7 may involve different neural circuits or states compared to the alpha-dominated seizures in other rats. Although we did not monitor or score sleep stages in this study, the dominance of delta-band power in some animals raises the possibility that seizure duration rhythmicity may be influenced by sleep–wake cycles. This represents an important direction for future work. Simultaneous monitoring of sleep stages and seizure dynamics would be valuable to determine whether seizure duration is modulated by specific sleep phases or circadian-dependent transitions

While we observed a consistent positive correlation between seizure duration and dominant EEG power in several rats, we interpret this relationship with caution. The increase in power may reflect the longer, more uniform rhythmic structure of prolonged seizures rather than a direct causal link. These findings may reflect co-modulation of both features by underlying network states. We also recognize that the specific frequency bands showing strongest associations varied across animals. This likely reflects inter-individual differences in dominant oscillatory activity, which are expected in both animal models and human studies. As such, we interpret these results primarily as intra-animal findings, highlighting that, within each subject, seizure duration may be modulated by fluctuations in dominant rhythms.

Several studies have examined the changes in power of the dominant frequency during ictal periods. For example, Bragin et al. (2005) [2] showed that in chronic seizure models using rats, 70% of seizures began with an increase in the frequency of interictal EEG spikes (hypersynchronous type), which evolved into low-voltage fast patterns [2]. Fisher et al. (1992) [1] demonstrated a significant increase in spectral power above 35 Hz during electrodecremental events at the start of seizures in patients with partial seizures, with power increasing up to fivefold in the 80-120 Hz portion of the spectrum [1]. In the study by Khosravani et al. (2009) [22], low-frequency oscillations (0–100 Hz) were found to increase in absolute magnitude during seizures. Furthermore, while high-frequency oscillations (HFOs) tend to be of low amplitude, they often override the larger, slower frequency waveforms that dominate the early phases of electrographic seizures, indicating a dynamic interplay between these frequency ranges during seizure activity [22]. While these studies provide valuable insight into the changes in dominant frequency power during seizures, they do not directly explore the relationship between seizure duration and dominant EEG power. Our study focuses on this relationship, providing novel insights into how dominant EEG power might influence seizure length.

### Probing increases seizure likelihood and prolongs seizure duration in rats

We also considered the potential influence of the probing protocol on seizure dynamics. Probing was conducted at regular intervals (5-minute stimulation periods followed by 5-minute off periods) throughout the study. However, this stimulation was not phase-locked or synchronized to any internal biological rhythm such as circadian or multidien cycles. The primary rhythmic patterns identified in seizure duration occurred on substantially longer timescales (4-8 days), making it unlikely that the probing schedule directly imposed these multiday rhythms. Nonetheless, we recognize that repeated stimulation may modulate cortical excitability and could influence seizure occurrence or duration, particularly in susceptible phases of ongoing cycles. To account for this, we explicitly compared seizure durations during probing-on and probing-off periods (6) and found statistically significant differences in some rats. While this supports the idea that probing can interact with seizure dynamics, it does not explain the presence of long-timescale periodicity.

The results presented in Fig 6 highlight the effect of probing on seizure occurrence and duration. Across all six rats (T2–T7), there is a higher frequency of seizures during the probing on periods compared to the off periods, suggesting that the probing stimulus used in the experiments increased the likelihood of seizures. This observation aligns with the notion that external stimulation, even at low levels, can act as a trigger for seizure activity, possibly lowering the threshold for seizure initiation. Frauscher et al. (2023) emphasized that stimulation can significantly impact brain dynamics, triggering seizures in some cases to aid in seizure onset localization and to provide insight into epileptic network connectivity [12]. It is important to note that probing effects are parameter-specific and not generalizable to all forms of stimulation.

Additionally, our results indicate significant differences in seizure durations between probing-on and probing-off periods in several rats, particularly in T2, T3, T4, and T7, with longer seizure durations observed during the probing-on periods (*p <* 0.05). This suggests that probing not only increases seizure frequency but may also enhance seizure intensity, potentially through the stimulation’s cumulative effect on neural activity.

### Clinical implications

Understanding the cyclic patterns in seizure duration offers valuable insights into the underlying neurophysiological sources of epilepsy and provides a basis for designing targeted interventions to control seizure severity. The discovery of rhythmic fluctuations in seizure duration suggests that epileptic activity is not purely random but may be influenced by intrinsic or extrinsic cycles, potentially linked to circadian rhythms, neurochemical fluctuations, or underlying neural network dynamics. By identifying these cycles, we can gain a clearer picture of the factors that drive seizure activity and develop strategies to mitigate its impact.

The presence of rhythmicity in seizure duration could indicate specific oscillatory processes within the brain that contribute to seizure generation and propagation. For instance, if seizure duration follows predictable patterns tied to the brain’s natural cycles, such as sleep-wake or hormone cycles, then interventions could be timed to periods of increased vulnerability or reduced threshold for prolonged seizures. This approach could help in developing therapies that are in sync with these cycles, either by introducing medication, neurostimulation, or other therapeutic techniques precisely when they are most needed or when the brain is most responsive to intervention.

Additionally, by understanding seizure duration cycles, clinicians and researchers could explore more deeply the structural or functional network changes that influence these patterns. For example, prolonged seizures that follow particular cycles might be connected to heightened activity in specific neural circuits or regions. Targeting these circuits with individualized treatments, such as responsive neurostimulation, could help regulate seizure duration more effectively. Further, recognizing cycles in seizure duration allows for a proactive, rather than reactive, approach to epilepsy management, enabling personalized care strategies that reduce the intensity or duration of seizures based on anticipated changes in cycle phases.

### Future directions

We acknowledge that this study was conducted exclusively in male rats, which limits the generalizability of our findings. As such, it remains an open question whether similar rhythmic patterns in seizure duration would be observed in female rats. Future studies should incorporate both sexes to better understand potential sex-specific dynamics in seizure evolution.

An exciting direction for future research involves the use of computational models to understand the neural circuitry underlying the observed cycles in both seizure frequency and duration. By simulating neural networks and brain circuits, computational models can help us explore the specific mechanisms that contribute to cyclic patterns in seizure activity. Such models could provide a virtual testing ground for hypotheses about the oscillatory behavior of neural circuits, enabling researchers to investigate how intrinsic rhythms, connectivity patterns, and external influences interact to shape seizure dynamics [9, 10, 25, 26].

Computational models could incorporate external factors, such as circadian or hormonal cycles, to simulate how these systems might interact with brain circuits to influence seizure cycles. This could help us understand why certain times of day or physiological states are associated with increased seizure frequency or duration and guide the timing of interventions to align with the brain’s natural rhythms. Moreover, models that simulate the impact of periodic or rhythmic interventions—such as targeted neurostimulation delivered in sync with identified cycles—could reveal optimized stimulation protocols aimed at modulating or disrupting seizure patterns more effectively. By advancing our understanding of these neural mechanisms, computational modeling could accelerate the development of innovative, rhythm-aware treatments aimed at mitigating seizure frequency and duration through targeted modulation of the underlying circuits. As these models become increasingly sophisticated and data-driven, they hold promise for transforming our approach to epilepsy management, paving the way for more effective, personalized care.

## Conclusion

This study provides novel insights into the cyclical patterns governing seizure duration, emphasizing the presence of significant rhythmic fluctuations in seizure dynamics. Through detailed analysis of six rats over a 40-day recording period, we identified that seizure duration follows predictable temporal structures, with distinct periodicities varying across individuals. Additionally, this work explored the influence of inter-seizure intervals and dominant EEG power, offering a broader perspective on the factors that may modulate seizure severity. The finding that seizure durations align with specific phases within identified cycles, supported by Synchronization Index (SI) analysis, reinforces the notion that seizure activity is governed by underlying temporal dynamics rather than occurring randomly. These insights hold potential clinical implications, suggesting avenues for time-based therapeutic interventions and personalized epilepsy management.

Our results highlight that understanding rhythmic patterns in seizure duration could aid in identifying neural mechanisms that contribute to epileptic activity and influence seizure characteristics. By establishing links between seizure duration and specific EEG frequencies, as well as exploring the impact of probing, this study contributes to a growing understanding of seizure modulation in epilepsy research. Future work, including computational modeling of seizure cycles, may further clarify the neural circuitry behind these rhythms, leading to targeted approaches that mitigate seizure severity through informed, rhythm-aware interventions.

## Acknowledgments

The authors thank Dr Warwick Cheung for providing the rat recordings. This work was funded by the Australian Government under the Australian Research Council’s Training Centre in Cognitive Computing for Medical Technologies (project number ICI70200030).

## Supporting information

**Fig S1.**
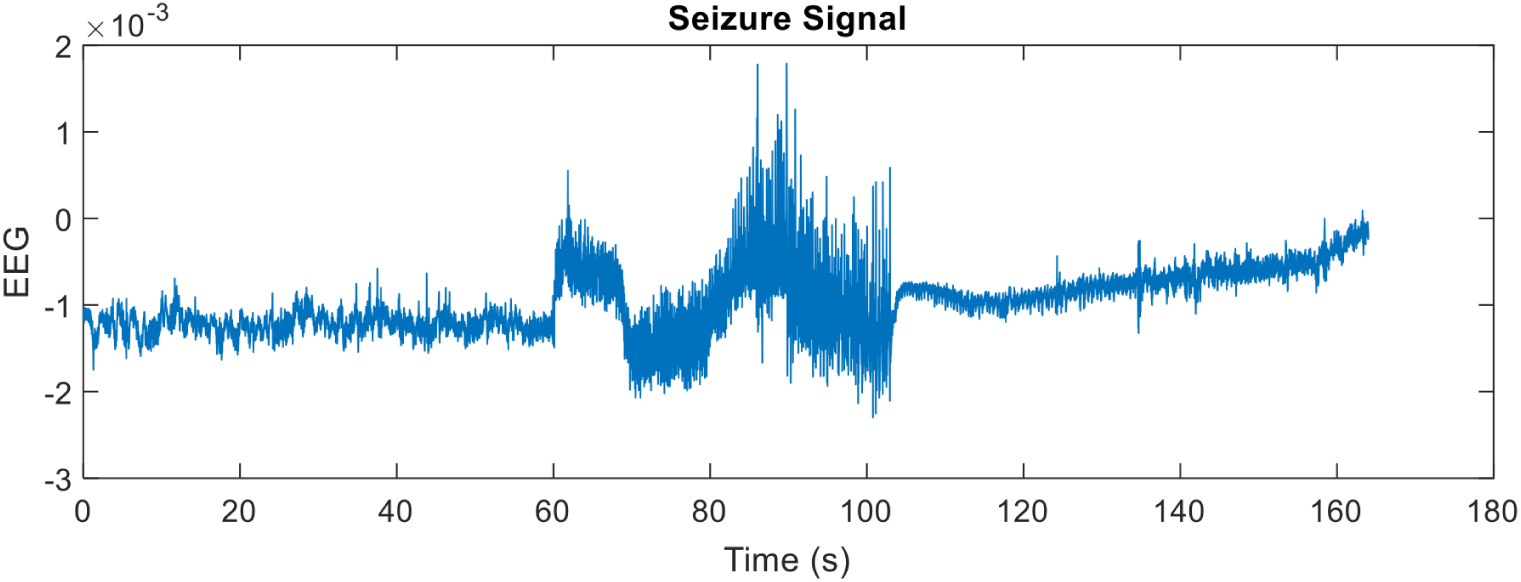
Representative EEG trace from the epilepsy model group, demonstrating the presence of a spontaneous seizure event.

**Fig S2.**
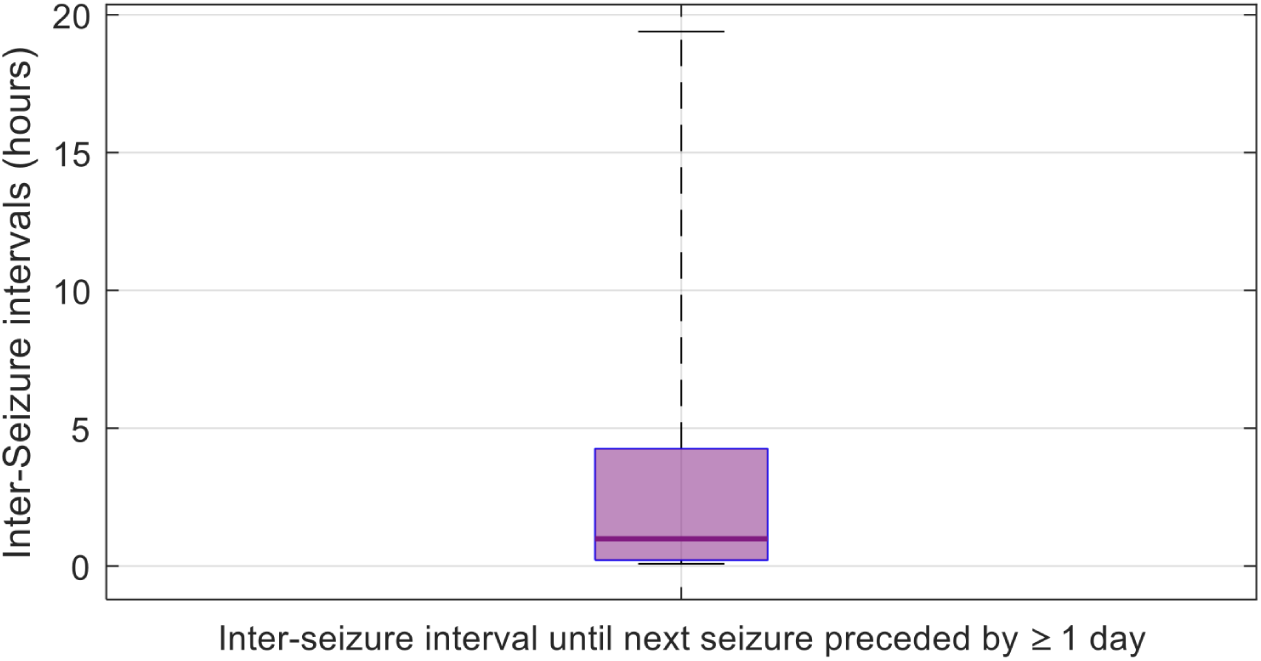
Inter-seizure intervals following seizures that were preceded by *≥*1 day of seizure-free time. The majority of these subsequent seizures occurred within a few hours, suggesting that seizures following long intervals are frequently followed by shorter-than-average recurrence times.

